# Microglia regulate motor neuron plasticity via reciprocal fractalkine/adenosine signaling

**DOI:** 10.1101/2024.05.07.592939

**Authors:** Alexandria B. Marciante, Arash Tadjalli, Kayla A. Burrowes, Jose R. Oberto, Edward K. Luca, Yasin B. Seven, Maria Nikodemova, Jyoti J. Watters, Tracy L. Baker, Gordon S. Mitchell

## Abstract

Microglia are innate CNS immune cells that play key roles in supporting key CNS functions including brain plasticity. We now report a previously unknown role for microglia in regulating neuroplasticity within spinal phrenic motor neurons, the neurons driving diaphragm contractions and breathing. We demonstrate that microglia regulate phrenic long-term facilitation (pLTF), a form of respiratory memory lasting hours after repetitive exposures to brief periods of low oxygen (acute intermittent hypoxia; AIH) via neuronal/microglial fractalkine signaling. AIH-induced pLTF is regulated by the balance between competing intracellular signaling cascades initiated by serotonin vs adenosine, respectively. Although brainstem raphe neurons release the relevant serotonin, the cellular source of adenosine is unknown. We tested a model in which hypoxia initiates fractalkine signaling between phrenic motor neurons and nearby microglia that triggers extracellular adenosine accumulation. With moderate AIH, phrenic motor neuron adenosine 2A receptor activation undermines serotonin-dominant pLTF; in contrast, severe AIH drives pLTF by a unique, adenosine-dominant mechanism. Phrenic motor neuron fractalkine knockdown, cervical spinal fractalkine receptor inhibition on nearby microglia, and microglial depletion enhance serotonin-dominant pLTF with moderate AIH but suppress adenosine-dominant pLTF with severe AIH. Thus, microglia play novel functions in the healthy spinal cord, regulating hypoxia-induced neuroplasticity within the motor neurons responsible for breathing.

## MAIN

Microglia are resident immune cells of the central nervous system. In the healthy brain, they also perform important physiological roles in neuronal homeostasis and regulation of synaptic plasticity by engaging in reciprocal communication with other cells through multiple ligands and receptors. For example, the neuro-chemokine fractalkine (Fkn; CX3CL1) is released from neurons in response to physiological stimuli such as neuronal activity or hypoxia, activating Fkn receptors on nearby microglia^1–3^. In the CNS, Fkn is expressed predominantly by neurons, whereas its receptor (CX3CR1) is expressed uniquely by microglia^4–6^. Previous studies indicate neuron-microglial Fkn signaling regulates important neural functions such as hippocampal synaptic plasticity^7,8^ and medullary respiratory rhythm generation^9^. Fractalkine modulates activity-dependent synaptic transmission *via* factors such as microglia-dependent increases in extracellular ATP and/or adenosine^3,10,11^. However, the role of microglia in regulating any form of spinal, motor or activity-independent synaptic plasticity is unknown.

The neural system controlling breathing exhibits considerable neuroplasticity^12^. Spontaneous respiratory plasticity is of considerable importance when confronted with intrinsic or extrinsic physiological conditions (*e.g*., weight gain or loss), or with the onset of clinical disorders that compromise breathing, such as neurological and lung/chest wall disorders^12,13^. A powerful stimulus to respiratory plasticity is intermittent hypoxia, or intermittent exposure to low oxygen. Intermittent hypoxia elicits plasticity in the oxygen-sensitive carotid body chemoreceptors^14,15^, their brainstem integrating neurons in the nucleus of the solitary tract^16^ or neurons critically involved in respiratory rhythm generation^17–19^. Intermittent hypoxia also elicits plasticity in the output neurons of the respiratory control system—respiratory (phrenic) motor neurons^20,21^. The phrenic motor system is located at cervical spinal segments 3-6, which provides innervation of the major inspiratory pump muscle, the diaphragm^22,23,24^.

The most extensively studied model of phrenic motor plasticity is known as phrenic long-term facilitation (pLTF), a persistent elevation in phrenic nerve activity lasting hours after exposure to 3, 5-minute episodes of moderate hypoxia (acute intermittent hypoxia; AIH). AIH elicits pLTF *via* distinct (but competing) cellular mechanisms, depending on the severity of hypoxic episodes within the AIH protocol^25^. Whereas AIH consisting of moderate hypoxic episodes (mAIH) elicits pLTF by a serotonin-dependent, adenosine-constrained mechanism, AIH consisting of severe hypoxic episodes (sAIH) elicits phenotypically similar pLTF by a distinct adenosine-driven and serotonin-constrained mechanism^25–28^. With intermediate hypoxia^29^ and pharmacological co-activation of the relevant cervical spinal serotonin (5-HT_2_) and adenosine (A_2A_) receptors, pLTF is canceled *via* powerful crosstalk inhibition^30^. Thus, the serotonin *versus* adenosine balance is a key regulator of AIH-induced phrenic motor plasticity.

While the relevant spinal serotonin is released from descending projections of brainstem raphe serotonergic neurons^27^, the cellular source of adenosine is unknown. Here we tested the hypothesis that AIH triggers phrenic motor neuron-microglial interactions mediated by fractalkine signaling that regulate the serotonin/adenosine balance and, thus, phrenic motor plasticity. Specifically, we tested the idea that, during hypoxic episodes, phrenic motor neuron Fkn activates microglial Fkn receptors (CX3CR1)—the lone cell type known to express CX3CR1 in the healthy CNS. In response, microglia increase conversion of ATP to adenosine, increasing extracellular adenosine concentration. Thus, phrenic motor neuron-microglia interactions regulate the local serotonin/adenosine balance *via* reciprocal Fkn/CX3CR1-adenosine signaling. We demonstrate that: 1) cervical spinal Fkn is sufficient to drive phrenic motor plasticity by evoking local ATP/adenosine release/production; 2) microglia constrain mAIH-induced pLTF due to adenosine accumulation, or (with sufficient adenosine accumulation) drive sAIH-induced, adenosine-dominant pLTF; 3) cervical spinal microglial CX3CR1 activation is necessary for sAIH-induced pLTF, but constrains mAIH-induced pLTF; and 4) the relevant Fkn protein is within phrenic motor neurons *per se*. Thus, through interactions with nearby microglia, phrenic motor neurons indirectly regulate the magnitude and mechanism of their own plasticity.

## RESULTS

### Fkn increases extracellular adenosine levels and elicits phrenic motor facilitation

We first tested whether spinal Fkn receptor activation near the phrenic motor nucleus is sufficient to increase spinal extracellular adenosine levels. Fkn protein was delivered *via* intrathecal injection to the C4 cervical spinal cord segment in anesthetized, paralyzed and ventilated rats; changes in cervical spinal extracellular adenosine concentration were measured via adenosine and inosine microbiosensors (**Figure 1A**). Phrenic nerve activity increased in a dose-dependent manner in response to intrathecal Fkn injections (*see* ***Supplementary Figure 1***; for details concerning experimental preparation or adenosine measurements, see *Methods* or^31–33^). Intrathecal Fkn injections (100 ng) slowly increased *in vivo* spinal extracellular adenosine concentration over ∼30 minutes (p < 0.001; **Figure 1B**). Concurrent blood gas measurements verified that rats remained well oxygenated (arterial PO_2_ = 291 ± 13 mmHg) and isocapnic (arterial PCO_2_ = 44.8 ± 0.5 mmHg) during recordings.

**Figure 1.**
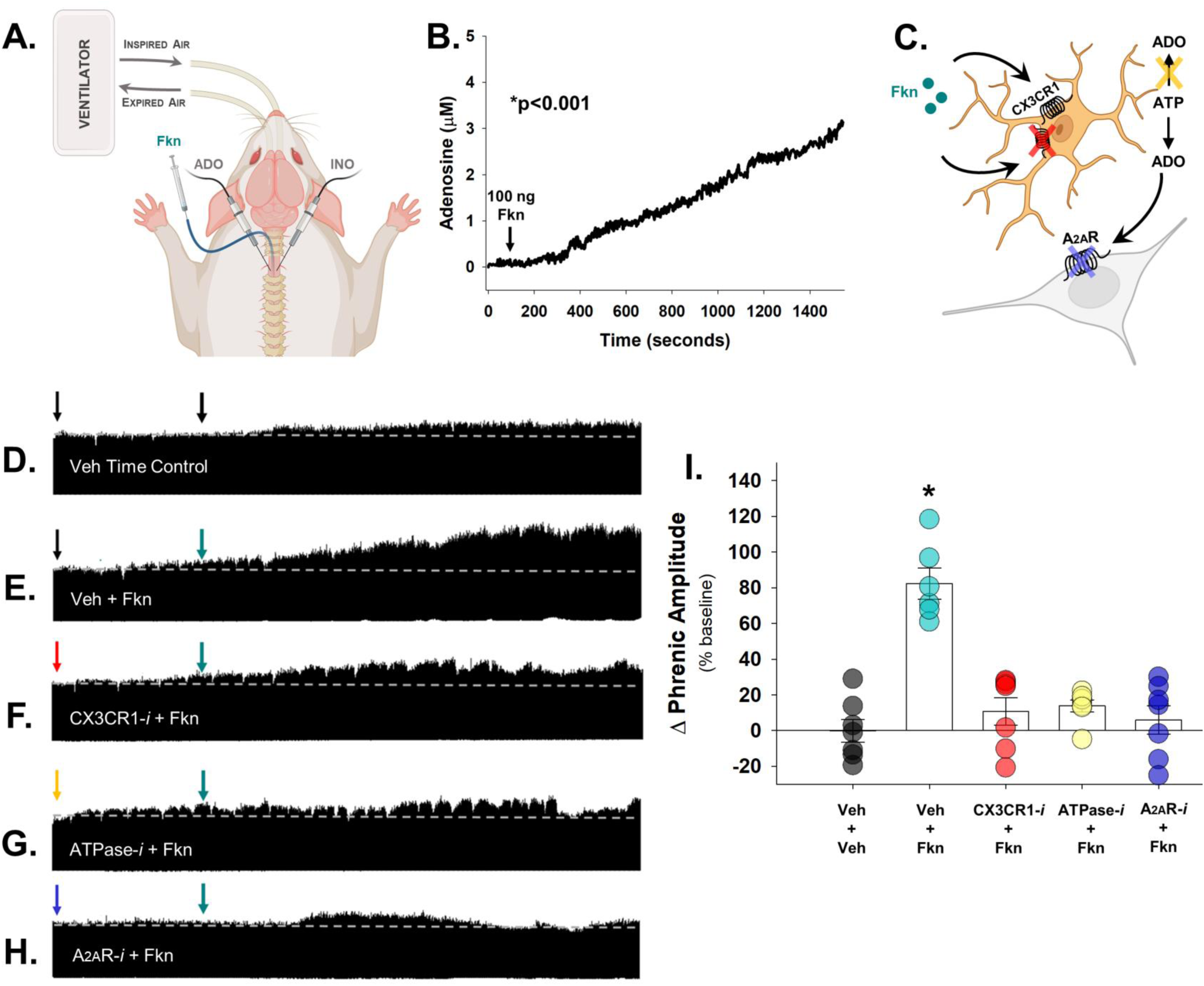
Fkn increases extracellular adenosine levels in the cervical spinal cord and elicits adenosine-dependent phrenic motor facilitation. (**A**): Schematic of experimental setup with intrathecal drug delivery and placement of adenosine (ADO) and inosine (INO) micro-biosensors. (**B**) Intrathecal fractalkine (Fkn; 100 ng; 12 μL) delivery evoked a slow increase in extracellular adenosine concentration over 30 min (n = 3 independent recordings; Linear Regression, p < 0.001; r = 0.993, r^2^ = 0.985, Adjusted r^2^ = 0.985). Concurrently, phrenic nerve activity was recorded in urethane anesthetized rats maintained at baseline conditions during and after intrathecal Fkn injection (90 min post-delivery). Ventilator volumes were set for each rat based on body mass (0.007 ml * body weight, g; 72-74 breaths per minute). Inspired CO2 or ventilator frequency were adjusted to maintain end-tidal PCO_2_ between 38 and 41 mmHg. Blood gas measurements were taken 2-3 times during the initial baseline, and at 30, 60, and 90-min post-drug (***Supplementary Table 1***). (**C**) Schematic of inter-cellular signaling highlighting where receptors/enzymes were blocked/inhibited in **D**-**H.** Representative compressed neurograms of integrated phrenic nerve activity are shown for rats that received (**D**) vehicle (VEH; time controls), (**E**) VEH + Fkn, (**F**) CX3CR1 inhibitor, AZD8797 + Fkn, (**G**) ATPase inhibitor, ARL67156 + Fkn, and (**H**) A2A Receptor inhibitor, MSX-3 + Fkn. (**I**) Phrenic burst amplitude (percent change from baseline; % baseline) was significantly increased 90 min post-Fkn administration (VEH + Fkn); however, CX3CR1, ATPase and A2A receptor inhibition (schematized in **C**) attenuated or prevented phrenic motor facilitation (n=6-7 recordings per group; F(4,29) = 21.378, p < 0.001; one-way ANOVA). ****p < 0.001, significant differences vs all groups; Tukey *post-hoc* Test. Bars show mean ± SEM.

To determine if Fkn-induced increases in extracellular adenosine concentration elicit phrenic motor facilitation, phrenic nerve activity was recorded in the same anesthetized rats (for details concerning experimental preparation, see *Methods*^31–33^). At 90 min post-injection, intrathecal Fkn significantly increased phrenic nerve burst amplitude (82.4 ± 8.7% above baseline *vs* vehicle time controls (VEH; −0.2 ± 6.4%; p<0.001; n=6-7 independent recordings per group; **Figures 1D-E**). Thus, Fkn receptor activation is sufficient to elicit phrenic motor facilitation.

We next tested the involvement of spinal receptors and enzymes in phrenic motor neuron-microglia Fkn signaling to assess their role in fractalkine-induced phrenic motor facilitation (**Figure 1C**). To this end, Fkn (100 ng) was injected over the phrenic motor pool 30 minutes after pretreatment with either cervical spinal: 1) microglial CX3CR1 inhibitor (**Figure 1F**); 2) ectonucleotidase inhibition to determine if extracellular ATP is converted to adenosine (*vs* adenosine release *per se*^34^ **Figure 1G**); and 3) A_2A_ receptor inhibition, a receptor necessary and sufficient for adenosine-driven phrenic motor plasticity^35^ (**Figure 1H**).

Each inhibitor prevented development of significant phrenic motor facilitation 90 min post-Fkn delivery (<u>CX3CR1 inhibition</u>: 10.7 ± 7.8%; <u>ATPase inhibition</u>: 13.7 ± 3.3%; <u>A_2A_ receptor inhibition</u>: 5.9 ± 7.9%; all p < 0.001 *vs* VEH + Fkn; **Figures 1I**). Thus, Fkn receptor activation on cervical microglia is sufficient to elicit spinal neuroplasticity *via* a mechanism that requires conversion of ATP to adenosine with subsequent A_2A_ receptor activation.

### Spinal extracellular adenosine accumulation is dependent on hypoxia severity

Within 5-minute hypoxic episodes, arterial hypoxemia invades cervical spinal tissues, lowering tissue PO_2_ to an extent that depends on the severity and duration of hypoxemia^36^. We now demonstrate that the severity of hypoxemia within episodes (and tissue hypoxia) has a major impact on spinal adenosine accumulation (*i.e*., mAIH *vs* sAIH). Hypoxia-evoked extracellular adenosine accumulation in the ventral cervical spinal cord near the phrenic motor nucleus exhibited dose-dependent adenosine accumulation during moderate (PaO_2_ = 43 ± 2 mmHg) *vs* severe (PaO_2_ = 27 ± 0.8 mmHg) hypoxic episodes (n = 5 recordings per group made in 3 rats; **Figures 2A** and **2B**). Peak adenosine levels were significantly higher during severe *vs* moderate hypoxic episodes (severe: 7.4 ± 0.8 µM; moderate: 3.3 ± 0.2 µM; t(8) = −5.299, p < 0.001; unpaired *t*-test; **Figure 2C**); when expressed as an area under the curve to reflect total adenosine accumulation during hypoxic episodes, adenosine accumulation during severe *vs* moderate hypoxic episodes remained elevated (severe: 32.2 ± 3.8 µM; moderate: 15.8 ± 0.7 µM; t(8) = −4.218, p = 0.003; unpaired *t*-test; **Figure 2D**). Thus, hypoxia evokes dose-dependent spinal adenosine accumulation near phrenic motor neurons (F(3,6)=229.816, p < 0.001; r = 0.9957, r^2^ = 0.9914; **Figure 2E**). Pretreatment with a CX3CR1 antagonist suppressed adenosine accumulation during severe hypoxia (***Supplemental Figure 2***), demonstrating that reciprocal fractalkine/adenosine signaling *via* microglial CX3CR1 activation is necessary for hypoxia-evoked adenosine accumulation.

**Figure 2.**
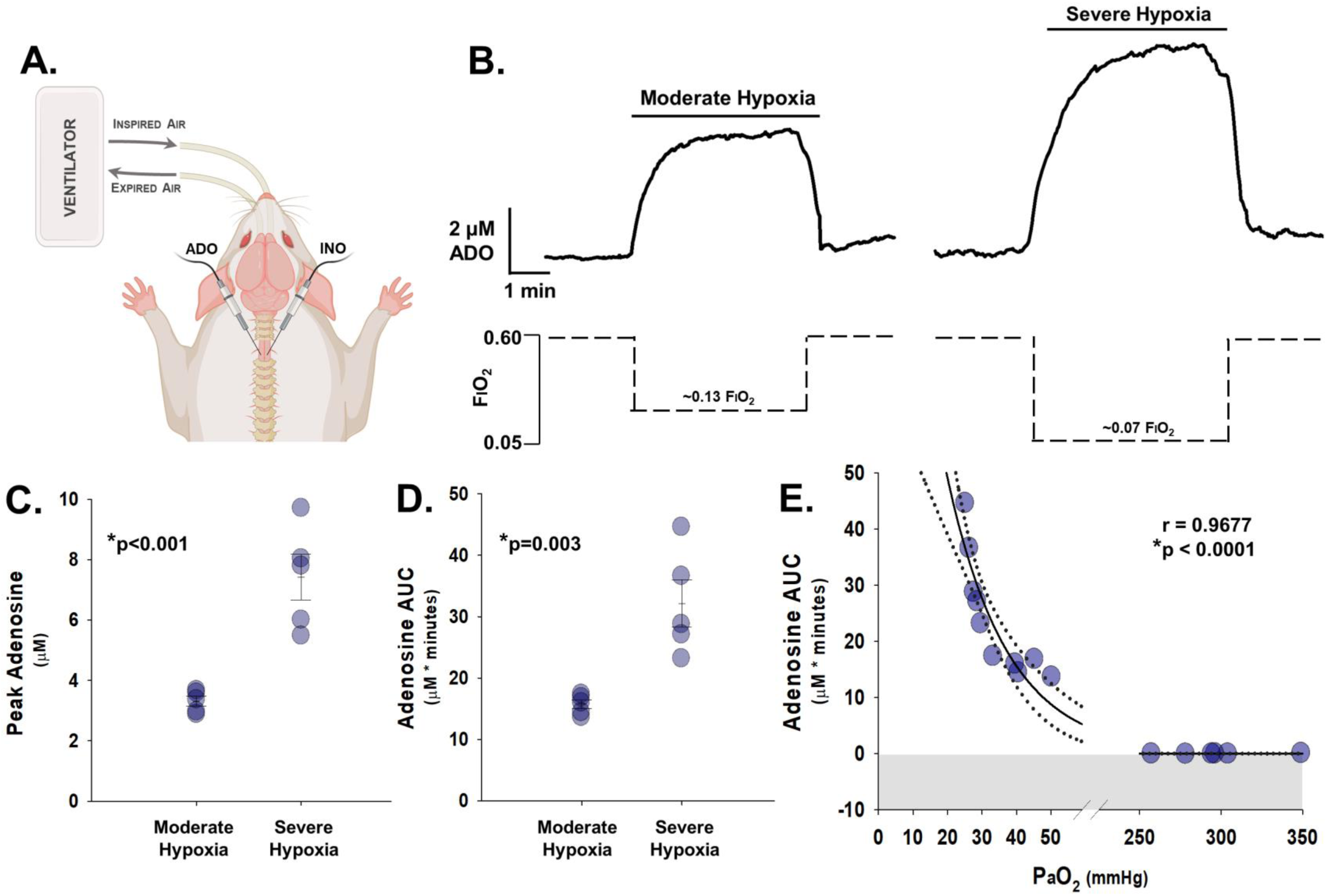
Severe (vs moderate) hypoxic episodes evoke greater spinal adenosine accumulation. (**A**) Adenosine/inosine probes placed between ventral C3/C4 to measure changes in adenosine accumulation during hypoxia. (**B**) Average traces of extracellular adenosine concentration (µM) during 5 min of moderate (PaO_2_ = 42.7 ± 2.0 mmHg) or severe (PaO_2_ = 27.2 ± 0.8 mmHg) hypoxia (n = 5 per group from 3 rats). Greater adenosine accumulation was observed in severe hypoxic episodes when expressed as (**C**) peak adenosine level ([ADO]peak; t(8) = −5.299, p < 0.001; unpaired *t*-test) or (**D**) total area under the curve ([ADO]AUC; t(8) = −4.218, p = 0.003; unpaired *t*-test). (**E**) PaO_2_ strongly correlates with measured extracellular adenosine levels (F(1,14)=206.099, p < 0.0001; r = 0.9677, r^2^ = 0.9364; Adjusted r^2^ = 0.9318 nonlinear regression). Bars are means ± SEM.

### Fractalkine signaling undermines moderate (serotonin-dominant) AIH-induced pLTF

Moderate AIH-induced phrenic long-term facilitation (pLTF) manifests as a persistent (and progressive) increase in integrated nerve burst amplitude lasting many hours after AIH has ended. The historical standard mAIH protocol^30^ consists of 3, 5-minute episodes of moderate hypoxia (arterial PO2 ∼40-55 mmHg), with 5 minute intervals of control oxygen conditions (arterial PO_2_ > 150 mmHg)^17,37,38^. To test the idea that Fkn/CX3CR1 signaling and adenosine accumulation during moderate AIH would suppress pLTF, intrathecal Fkn was delivered prior to moderate AIH (100 ng in 12 µL). As predicted, intrathecal fractalkine, delivered 30 minutes before abolished moderate AIH-induced pLTF (−1.7 ± 5.1%; *vs* vehicle controls: 43.2 ± 5.4%; p < 0.001; n=7-8 each group; **Figure 3A**). Fkn delivered prior to moderate AIH was unable to drive adenosine-dependent phrenic motor plasticity since concurrent serotonin and adenosine receptor activation cancels phrenic motor plasticity^30^.

**Figure 3.**
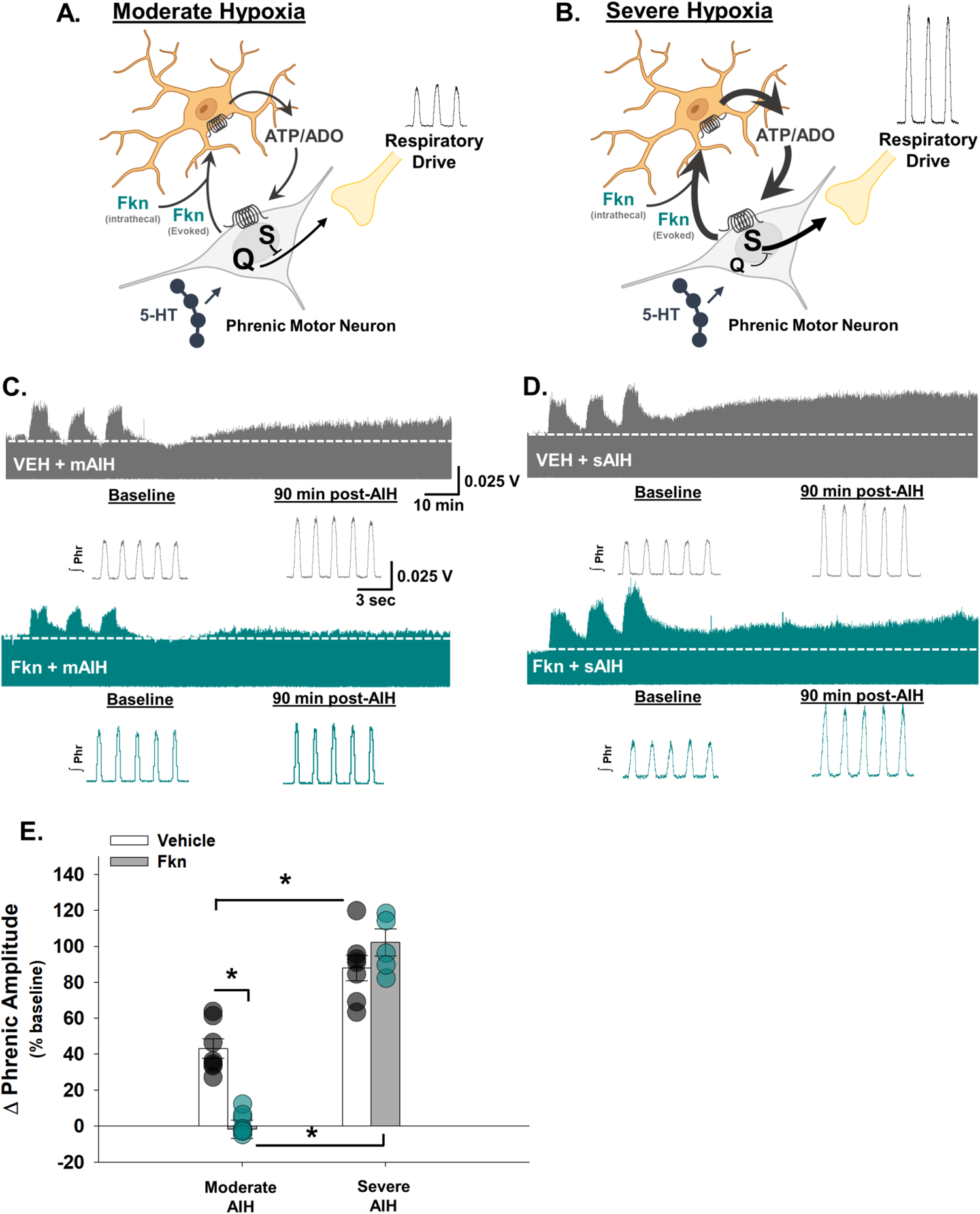
Intrathecal fractalkine differentially regulates moderate (serotonin-dominant) vs severe (adenosine-dominant) AIH-induced pLTF. Schematic of hypothesized mechanisms for intrathecal and hypoxia-evoked fractalkine (Fkn) release on moderate (**A:** serotonin-driven Q pathway) *vs* severe AIH-induced pLTF (**B:** adenosine-driven S pathway). Phrenic nerve activity was recorded in urethane anesthetized rats during baseline, during intrathecal drug administration, and for 90 min post-treatment while baseline conditions were maintained. Inspired CO2 and/or ventilator frequency was adjusted to maintain end-tidal PCO2 between 38 and 41 mmHg. Blood gas measurements were taken 2-3 times during the initial baseline, during the last minute of the first hypoxic episode, and at 30, 60, and 90 min post-AIH (***Supplementary Tables 1*** and ***2***). Raw integrated phrenic nerve amplitude at baseline and during maximal chemoreceptor stimulation (10% O2, 7% CO2, balance N2) delivered at the end of each experiment are included in ***Supplementary Table 3*** to assess recording quality. (**C**, **D**) Representative compressed neurograms of integrated phrenic nerve activity from rats that received vehicle (VEH; *top row*) or fractalkine (Fkn; *bottom row*) ∼30 min prior to moderate (**C**) or severe (**D**) AIH. Immediately below each neurogram are individual, integrated (**∫**) phrenic nerve bursts taken during baseline and 90 min post-AIH. One-minute averages of phrenic nerve amplitude were measured at 90 min post-AIH, and are presented as percent change from the pre-AIH baseline value (**E**); there was a statistically significant interaction between drug (VEH *vs* Fkn) and AIH protocol (moderate *vs* severe) on pLTF (n=5-8 independent recordings each group; F(1,23) = 22.316, p < 0.001; two-way ANOVA). *p < 0.001; Tukey *post-hoc* Test. Bars are means ± SEM.

Severe AIH consisting of 3, 5 min episodes of severe hypoxia (PaO_2_: 25-30 mmHg), with 5 min intervals of baseline oxygen conditions, elicits pLTF by an adenosine-dominant mechanism^21,25^. Thus, in striking contrast to moderate AIH, intrathecal Fkn does not significantly affect severe AIH-induced pLTF (102 ± 7%; n=5; *vs* vehicle controls: 88 ± 7%; n=7; p = 0.159; **Figure 3**), consistent with the idea that adding Fkn-evoked adenosine beyond to the already high degree of A_2A_ receptor activation during severe AIH reinforces rather than inhibits plasticity. Thus, the same treatment, Fkn, undermines serotonin-driven, but not adenosine-driven pLTF.

### Microglia differentially regulate moderate vs severe AIH-induced pLTF

Although multiple cell types potentially generate extracellular ATP/adenosine during hypoxia, the source of spinal adenosine relevant to pLTF is unknown. Here, we demonstrate that microglial ablation and/or CX3CR1 receptor inhibition on spinal microglia have similar effects on phrenic motor plasticity. First, cervical spinal CX3CR1 inhibition enhanced moderate (serotonin-dominant) but attenuated severe (adenosine-dominant) AIH-induced pLTF. A selective CX3CR1 inhibitor was delivered intrathecally ∼30 minutes prior to moderate (**Figure 4A**) or severe AIH (**Figure 4B**) to antagonize cervical spinal fractalkine receptors. While severe AIH-induced pLTF is typically more robust than moderate AIH-induced pLTF (**Figure 4C**; p < 0.001), cervical spinal CX3CR1 inhibition significantly increased moderate AIH-induced pLTF (82 ± 5%) *vs* vehicle controls (43 ± 5%; n=7 each group; p < 0.001), but abolished severe AIH-induced pLTF (1 ± 6%) *vs* vehicle controls (88 ± 7%; n=7 per group; p < 0.001 *vs* vehicle controls; **Figure 4C**). These findings are consistent with the idea that cervical spinal CX3CR1 inhibition reduced extracellular adenosine accumulation during hypoxic episodes (***Supplementary Figure 2***).

**Figure 4.**
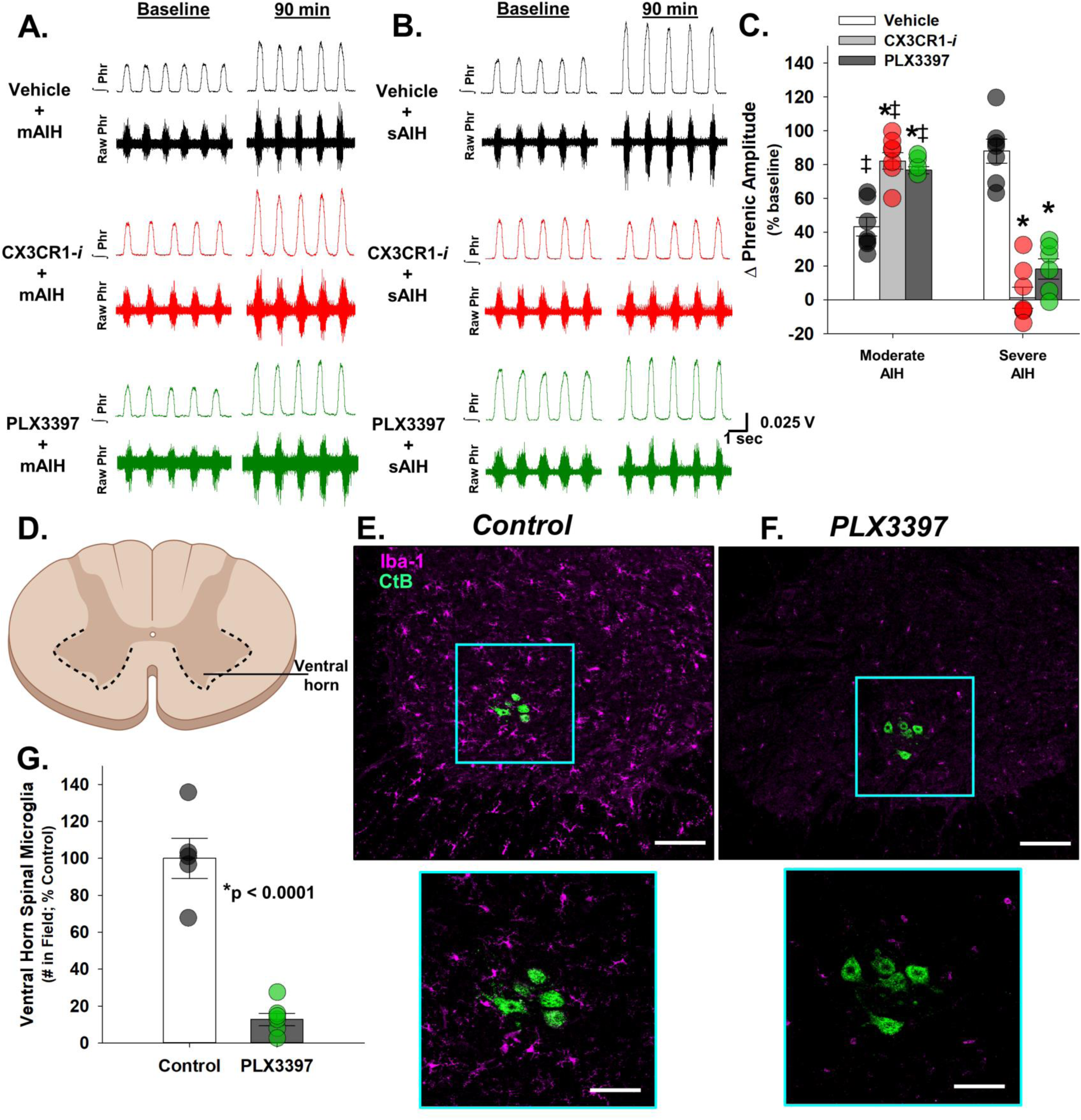
Microglia differentially regulate moderate (serotonin-driven) vs severe (adenosine-driven) AIH-induced pLTF. (**A**, **B**) Representative individual, integrated (**∫**) and raw phrenic (Phr) nerve bursts during baseline and 90 min post-AIH from a vehicle (VEH) control (*top row*), spinal CX3CR1-inhibited (CX3CR1*i*) rat, and PLX3397-treated rat (*bottom row*) prior to moderate (mAIH; **A**) or severe AIH (sAIH; **B**). One-minute averages of phrenic nerve burst amplitude were measured 90 min post-AIH, and are presented as percent change from baseline (**C**). There was a statistically significant interaction between drug (VEH *vs* CX3CR1*i or* PLX3397) and AIH protocol (moderate *vs* severe) on pLTF (n=4-7 independent recordings each group; F(2,32) = 69.896, p < 0.001; two-way ANOVA). *p ≤ 0.001, significant differences vs VEH controls; ‡p < 0.001, significant differences vs sAIH group pretreated with the same drug; Tukey *post-hoc* Test. (**D**) Schematic outlining region of the ventral horn where microglia were counted after VEH or PLEX3397 treatment. Representative confocal microscope images from VEH control (**E**) and PLX3397-treated rats (**F**) stained for Iba-1 positive microglia (magenta) and phrenic motor neurons (CtB; green). Scale bar (left; 10x magnification): 150 µm; scale bar (right; 40x magnification): 50 µm. (**G**) Ventral horn Iba-1 positive microglia were counted using a custom code; Iba-1 positive microglia counts were significantly reduced in spinal cords of PLX3397-treated rats *versus* VEH controls (spinal tissue from n=5-6 rats each group with at least 10 sections per rat; t(9) = 8.347, p = 0.00002; unpaired *t*-test). Bars are means ± SEM.

Several cell types (*e.g.*, microglia, astrocytes and neurons) could release ATP/adenosine during hypoxia and contribute to increased extracellular adenosine levels. To determine if microglia *per se* are necessary for the hypoxia-evoked adenosine increase (releasing adenosine or converting extracellular ATP from other cells to adenosine *via* microglial ectonucleotidases^1^), microglia were ablated by treating rats with a potent colony-stimulating factor 1 (CSF1) receptor inhibitor^39^. CSF1 receptor signaling is required for CNS microglial survival^40^. After 10 days of PLX3397 administration (80 mg/kg/day *via* syringe-feeding), microglial depletion was verified *via* computer-assisted counting of Iba-1 positive cells using a custom code in cervical ventral horn (C3-C6; the region containing the phrenic motor nucleus; **Figure 4D-F**). Iba-1 positive cells were counted only if they colocalized with nuclear DAPI staining. Indeed, Iba-1 positive microglia counts were significantly reduced in PLX3397-treated rats (to 13 ± 3% of control; n=5-6 per group; t(9) = 8.347, p = 0.00002; unpaired *t*-test; **Figure 4G**). Moderate (serotonin-dominant) AIH-induced pLTF increased nearly 2-fold in PLX3397-treated rats (77 ± 2%; n=4; *vs* vehicle controls, p = 0.001; **Figure 4C**). In contrast, severe (adenosine-dominant) AIH-induced pLTF was reduced 4-fold in PLX3397-treated rats (18 ± 6%; n=6; *vs* vehicle controls, p < 0.001; **Figure 4C**). Thus, microglial CX3CR1 inhibition and selective microglial ablation both shift the balance in favor of serotonin-*vs* adenosine-dominant plasticity.

### Phrenic motor neurons are the relevant source of fractalkine for phrenic motor plasticity

Fractalkine is abundantly expressed in CNS neurons and is released from those neurons during hypoxia^4,6,41,42^. While dorsal horn neurons are known to express Fkn^43,44^, reports of Fkn expression in alpha motor neurons are limited. Here, we demonstrate phrenic (and non-phrenic) motor neurons in the ventral cervical spinal cord express Fkn protein^45^; ***Supplementary Figure 3***). However, it is uncertain if the relevant fractalkine for microglial regulation of AIH-induced pLTF originates within phrenic motor neurons *per se vs* other neurons. Thus, small interfering RNAs (siRNA) targeting fractalkine mRNA (siFkn) or a non-targeting sequence (controls; siNTg) were delivered to phrenic motor neurons *via* intrapleural injections at the 5^th^ intercostal space along the anterior axial line on 3 consecutive days as described by others ^46–48^. Intrapleural siRNA pools with the Accel modification (Dharmacon) are taken up by phrenic motor neuron axon terminals and retrogradely transported to their cell bodies where they selectively knock-down target mRNA (and protein)^46–49^ (**Figure 5A**). One day after the final siRNA injection, rats were prepared for *in vivo* recordings of phrenic nerve activity and exposed to moderate or severe AIH. Phrenic motor neuron fractalkine knockdown *via* siRNAs significantly increased serotonin-dominant moderate AIH-induced pLTF (104 ± 7 %; n=7) *vs* siNTg controls (55 ± 5%; n=4; p = 0.002, Tukey *post-hoc* Test; **Figure 5B-D**), consistent with reduced adenosine constraint of serotonin-driven pLTF. In contrast, severe AIH-induced (adenosine-dominant) pLTF was attenuated by siFkn (22 ± 9%; n=7) *vs* siNTg controls (102 ± 16%; n=4; p < 0.001, Tukey *post-hoc* Test). There was a significant interaction between siRNA (siFkn or siNTg) and AIH protocol on pLTF expression (F(1,18) = 45.431, p < 0.001).

**Figure 5.**
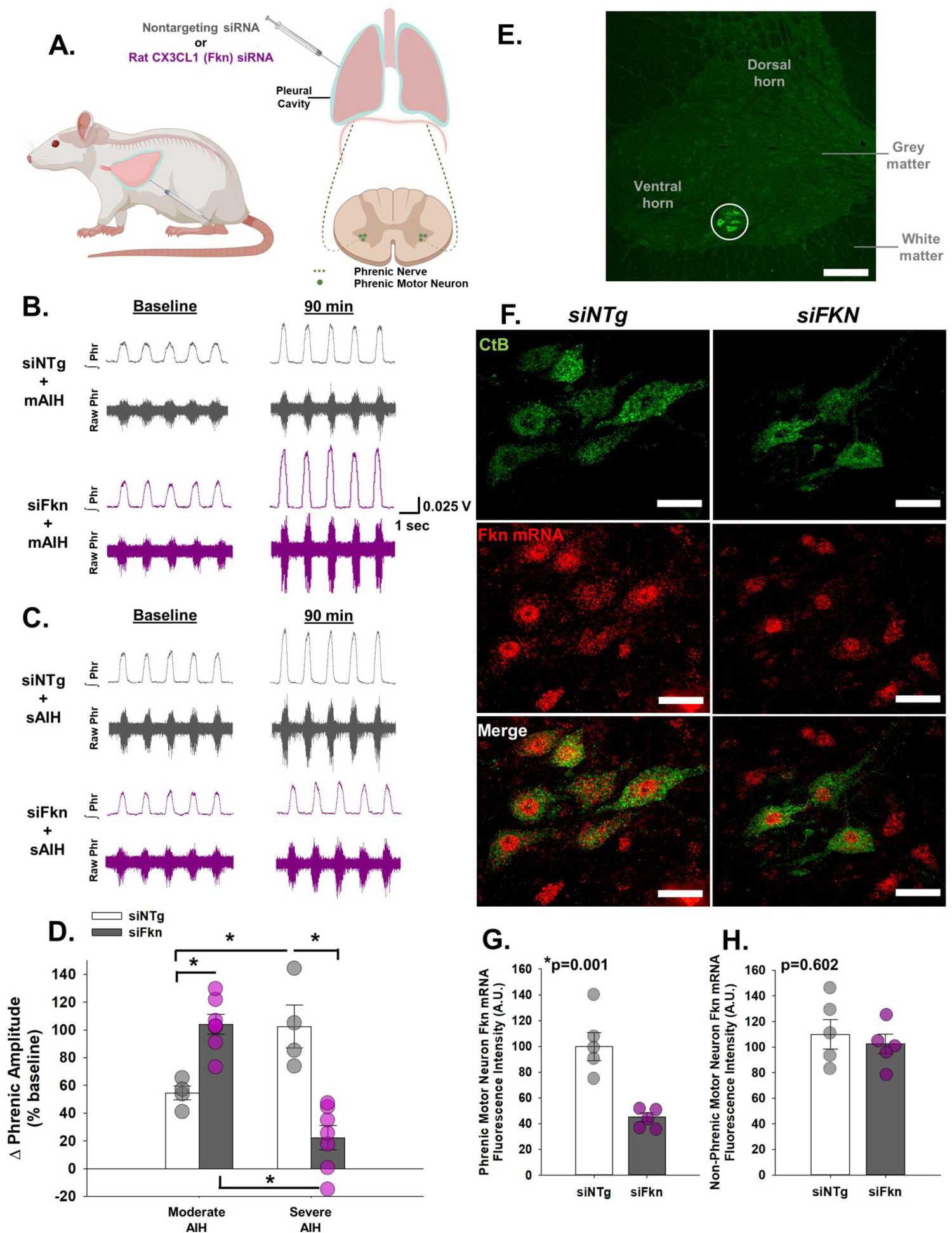
Phrenic motor neuron fractalkine undermines moderate (serotonin-dominant) AIH-induced pLTF but is required for severe (adenosine-dominant) AIH-induced pLTF. (**A**) Schematic depicting intrapleural injections for retrograde siRNA transport (nontargeting controls, siNTg; rat CX3CL1/fractalkine, siFkn) and CtB to phrenic motor neurons. (**B**, **C**) Representative individual, integrated (**∫**) and raw phrenic (Phr) nerve bursts taken during baseline and 90 min post-AIH from siNTg-(*top row*) and siFkn-injected rats (*bottom row*) prior to moderate (mAIH; **B**) or severe AIH (sAIH; **C**). One-minute averages of phrenic nerve amplitude were measured 90 min post-AIH, and are presented as percent change from baseline (**D**); there was a statistically significant interaction between siRNA (siNTg *versus* siFkn) and AIH protocol (moderate *vs* severe) on pLTF (n=4-7 independent recordings per group; F(1,18) = 45.431, p < 0.001; two-way ANOVA). *p < 0.01; Tukey *post-hoc* Test. (**E**) Confocal microscope image (10x magnification) of cervical ventral horn; phrenic motor nucleus circled; phrenic motor neurons identified with CtB (green); scale bar (10x magnification): 150 µm. (**F**) Representative confocal microscope images from siNTg (*left*) and siFkn (*right*)-injected groups stained for CtB (green) and Fkn mRNA (red); scale bar (40x magnification): 20 µm. (**G**) Fkn mRNA fluorescence intensity was significantly reduced in siFkn *vs* siNTg phrenic motor neurons (spinal tissue from n=5-6 rats per group with at least 10 sections per rat; t(8) = 4.812, p=0.001; unpaired *t*-test). (**H**) Fractalkine mRNA fluorescent intensity in non-phrenic motor neurons was similar between siNTg and siFkn groups (t(8) = 0.542, p=0.602; unpaired t-test). Bars are means ± SEM.

Phrenic motor neuron Fkn mRNA knockdown was verified after *in vivo* neurophysiology experiments had been completed *via* fluorescent *in situ hybridization*. Phrenic motor neurons were identified in the cervical spinal cord *via* intrapleural injections of cholera toxin B fragment (CtB) to retrogradely label phrenic motor neurons (**Figure 5E**). Abundant Fkn mRNA was observed in the nucleus and cytoplasm of (CtB-labeled) phrenic motor neuron somata (**Figure 5F**). Fkn mRNA was quantified within CtB-labeled phrenic motor neurons for siNTg and siFkn rats; fluorescence intensity at the phrenic motor nucleus was reduced nearly 60% in siFkn *vs* siNTg treated rats (t(8) = 4.812, p=0.001; unpaired *t*-test; **Figure 5G**). To determine if siFkn affected Fkn expression in other cells, fluorescent intensity was measured in nearby non-phrenic motor neurons, detected as large ventral horn cells (>300 μm^2^); no changes in fluorescence intensity could be appreciated in these cells (t(8) = 0.542, p=0.602; unpaired t-test; **Figure 5H**), consistent with prior reports of selective intrapleural siRNA delivery to phrenic motor neurons^46,48^. Collectively, these findings demonstrate that the relevant Fkn regulating AIH-induced phrenic motor plasticity is within phrenic motor neurons *per se*, demonstrating a means through which phrenic motor neurons regulate their own plasticity.

## DISCUSSION

We demonstrate that spinal microglia regulate the magnitude and mechanism of phrenic motor plasticity. We propose an intercellular model of reciprocal phrenic motor neuron-microglial interactions initiated by hypoxia-evoked phrenic motor neuron Fkn to microglial Fkn receptor signaling (**Figure 6**). During AIH, phrenic motor neuron Fkn (CX3CL1) activates Fkn receptors (CX3CR1) on nearby microglia, triggering conversion of extracellular ATP to adenosine. Depending on the severity of hypoxic episodes within the AIH protocol, hypoxia-evoked adenosine formation either constrains serotonin-driven pLTF or evokes pLTF by a distinct, adenosine-driven mechanism. Beyond demonstrating new roles for microglia in regulating hypoxia-evoked spinal respiratory motor plasticity^30^, these results have profound implications for ongoing translational efforts to harness AIH as a therapeutic modality to restore breathing and non-respiratory motor functions in severe clinical disorders that compromise breathing and other movements, such as spinal cord injury, ALS and multiple sclerosis^13,50–52^.

**Figure 6.**
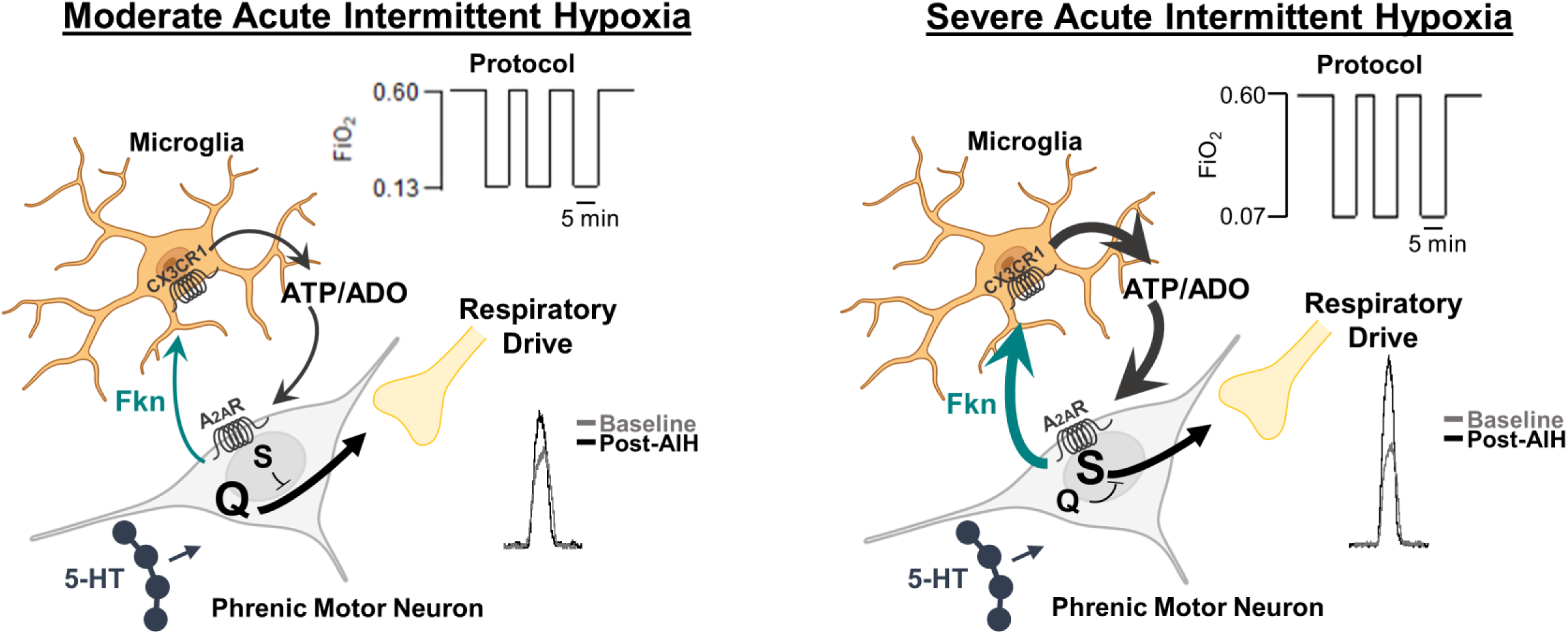
Intercellular model of reciprocal phrenic motor neuron Fkn to microglial CX3CR1 interactions during moderate and severe AIH-induced phrenic motor plasticity. Hypoxia triggers phrenic motor neuron Fkn signaling in a hypoxia dose-dependent manner. Fkn binds to its receptor, CX3CR1, on nearby microglia, triggering microglia-dependent formation and accumulation of extracellular adenosine (ADO). ADO activates phrenic motor neuron adenosine 2A (A_2A_) receptors, constraining the serotonin-dominant (Q) pathway to pLTF during moderate hypoxia (*left*). In contrast, severe hypoxia, and greater extracellular adenosine accumulation, shifts the serotonin/adenosine balance sufficiently to drive the adenosine-dominant S pathway to pLTF (*right*).

### Fkn-adenosine signaling between phrenic motor neurons and microglia regulate spinal neuroplasticity

In other regions of the central nervous system, there is growing awareness that neuron/glia interactions regulate activity-dependent synaptic plasticity. Both astroglia and microglia regulate brain plasticity in healthy animals^53–55^. We extend these principles to the spinal cord, specifically to the “final common pathway”—alpha motor neurons. Reciprocal Fkn-adenosine signaling between phrenic motor neurons and microglia has opposite effects on AIH-induced phrenic motor plasticity depending on a single variable: the severity of hypoxic episodes. Another important finding is that principles based on studies of hippocampal activity-dependent synaptic plasticity^56–58^ extend to <u>activity-independent</u>, serotonin-induced pLTF^30,59,60^. Collectively, the data presented here demonstrate that reciprocal phrenic motor neuron-microglia signaling has complex effects on activity-independent, AIH-induced phrenic motor plasticity, and that its impact depends on the severity of hypoxic episodes (**Figure 6**).

Perhaps the most significant finding of this study is that phrenic motor neurons are able to regulate their own plasticity *via* interactions with nearby microglia. To our knowledge, this is the first report that demonstrates *any* population of alpha motor neurons to interact with nearby microglia to regulate plasticity. Fkn knockdown within phrenic motor neurons demonstrates these are the neurons containing the relevant Fkn mRNA and protein. Although motor neuron-microglia interactions differentially impact plasticity elicited by moderate (suppression) *vs* severe AIH (activation), the apparent complexity can be explained by dose-dependent adenosine accumulation, biasing a system characterized by competing serotonin *vs* adenosine dependent mechanisms of plasticity^30^; each mechanism competes for dominance *via* powerful and mutual cross-talk inhibition. Even subtle shifts in the adenosine/serotonin balance can profoundly impact the magnitude and mechanism of plasticity^20,30,33^.

Moderate hypoxia rapidly activates carotid chemo-afferent neurons that project to and activate brainstem circuits, including raphe serotonergic neurons that project to the phrenic motor nucleus^61^. There, serotonin type 2 receptor activation initiates intracellular signaling cascades giving rise to pLTF^27,32,60^. Spinal tissue hypoxia evolves more slowly during hypoxic episodes of the AIH protocol^31,62^ and evokes the extracellular adenosine accumulation^33^ that constrains serotonin-driven plasticity *via* phrenic motor neuron A_2A_ receptor activation^27,63^. The intracellular signaling cascades initiated by Gq-coupled serotonin 2 *vs* Gs-coupled A_2A_ receptors are distinct^20,21^, other than their crosstalk interactions^28,49,64^. Thus, by shifting the serotonin/adenosine balance, microglia regulate the magnitude and mechanism of pLTF^20,25,35,65^. Whereas modest adenosine accumulation during moderate hypoxia constrains serotonin-dominant pLTF, more adenosine accumulates during severe hypoxic episodes, tipping the balance towards adenosine-dominant plasticity^25,46,66^. As a consequence, cervical spinal A_2A_ receptor inhibition enhances moderate AIH-^26^ but virtually blocks severe AIH-induced pLTF^25^. The present study demonstrates these relationships result from reciprocal phrenic motor neuron/microglial Fkn/adenosine signaling within the phrenic motor nucleus. Of considerable importance, the source of relevant Fkn mRNA and protein is the phrenic motor neurons (**Figure 5**).

### Spinal Fkn receptor activation modulates local extracellular adenosine levels

In the central nervous system, Fkn is synthesized predominantly by neurons, whereas its receptor (CX3CR1) is expressed exclusively by microglia^4–6^. By activating microglial Fkn receptors, Fkn initiates the neuron-microglia communication that: 1) coordinates inflammatory responses and suppresses overproduction of microglial pro-inflammatory factors^67,68^; 2) stimulates microglial migration towards neurons^6,69,70^; 3) regulates hippocampal synaptic plasticity^7,8^; and 4) modulates breathing^9^. We extend these concepts by demonstrating Fkn regulates spinal cord extracellular adenosine levels.

Spinal CX3CR1 inhibition blunts hypoxia-evoked extracellular adenosine accumulation (***Supplemental Figure 2***) as well as sAIH-induced adenosine-dominant pLTF and enhances mAIH-induced, serotonin-dominant pLTF (**Figure 3**). To further demonstrate microglial cells themselves regulate AIH-induced plasticity, microglial ablation with the CSF1 receptor inhibitor PLX3397 mimics Fkn receptor inhibition. Thus, microglia *per se* mediate bidirectional Fkn/CX3CR1 regulation of phrenic motor plasticity in moderate *vs* severe hypoxia, consistent with their postulated role as “adenosine donors” that constrain (mAIH) or enhance (sAIH) pLTF in a dose-dependent manner.

In the microglial BV2 cell line, adenosine accumulates in the media following Fkn receptor activation^2^, consistent with our hypothesis that microglia convert ATP and adenosine in response to Fkn receptor activation via microglial ectonucleotidases. Microglia express 2 membrane-bound ectonucleotidases: CD39 (converts ATP to ADP or AMP) and CD73 (converts AMP to adenosine^71^). We verified that cervical spinal microglia express CX3CR1, CD39 and CD73 (data not shown), and Fkn-induced that phrenic motor plasticity is impaired by an ectoATPase inhibitor, suggesting increased adenosine results from microglial ATPase activity (**Figure 1**). However, the specific cells contributing to extracellular ATP accumulation is unclear.

### Fkn within phrenic motor neurons regulates phrenic motor plasticity

One of the most compelling findings of our study is that RNA interference and Fkn knockdown within phrenic motor neurons exclusively impacts mAIH and sAIH-induced pLTF in a manner similar to spinal CX3CR1 inhibition, A_2A_ receptor inhibition and microglial ablation. Although non-phrenic motor neurons and interneurons also express Fkn, Fkn mRNA and (presumably) protein knockdown within phrenic motor neurons: 1) enhances serotonin-dominant, mAIH-induced pLTF; and 2) attenuates adenosine-dominant, sAIH-induced pLTF. Since rats have only ∼500 to 600 phrenic motor neurons amidst >3 million cells in C_3_-C_5_ spinal segments^23,72,73^, the impact of phrenic motor neuron Fkn knock-down is striking, and demonstrates that these neurons play a key role in regulating their own plasticity.

### Microglia are key regulators of extracellular adenosine levels

Fractalkine/CX3CR1 signaling is an important regulator of hippocampal synaptic plasticity^2,3,74,75^. Differential microglial expression of pro-inflammatory cytokines (IL6, TNFα, IL1β), as well as BDNF in ventral *vs* dorsal hippocampus, is associated with reduced dorsal and increased ventral LTP^74^, suggesting complexity similar to our findings with spinal, phrenic motor plasticity. With impaired CX3CR1 signaling, hippocampal LTP can be restored by IL1β receptor antagonists^76^. Whereas these studies suggest microglial <u>cytokines</u> may mediate Fkn effects on activity-dependent hippocampal LTP, reciprocal motor neuron/microglia signaling requires Fkn-induced <u>adenosine accumulation</u>.

We previously reported that mild systemic inflammation abolishes mAIH-induced pLTF by an adenosine-dependent mechanism^77^. However, since both astroglia and microglia mediate CNS pro-inflammatory responses^78–80^, the specific inflammation-activated cells undermining pLTF are not known. Our finding that healthy microglia are key regulators of phrenic motor plasticity, and that they act by favoring adenosine-dependent mechanisms to inhibit or drive plasticity suggests that mild systemic inflammation may also act *via* microglial activation^32^.

In conclusion, despite serving as CNS resident immune cells, brain microglia also play key roles in non-innate immune functions essential for normal CNS functioning. The present studies demonstrate that spinal microglia regulate plasticity in phrenic motor neurons *via* reciprocal Fkn/adenosine signaling between phrenic motor neurons and nearby microglia. This regulatory feed-back loop impacts plasticity in a hypoxia dose-dependent manner. Collectively, the findings reported here advance our fundamental understanding of phrenic motor plasticity, an activity-independent form of synaptic plasticity, is not cell-autonomous and requires interactions (incoming) raphe neurons, nearby microglia and phrenic motor neurons^30^. Understanding the regulation of AIH-induced phrenic motor plasticity may accelerate translation of AIH as a therapeutic modality to preserve/restore breathing (and non-respiratory movements) in devastating neurological conditions that end life due to respiratory failure, including cervical spinal cord injury, Amyotrophic Lateral Sclerosis and other disorders such as multiple sclerosis, stroke or traumatic brain injury^13^.

## METHODS

### Animals

All experiments were approved by the University of Florida Institutional Animal Care and Use Committee. Experiments were performed on young adult (3-4 months; 395 ± 5 g) male Sprague-Dawley rats (208A Colony, Envigo; IN, USA). Rats were housed in pairs at 24°C with a 12/12 light/dark cycle (lights on: 07:00; lights off: 19:00) with access to food and water *ad libitum*. All rats underwent a 14-day acclimation period prior to experiments. Sample size estimation was based on power analysis of previous studies^26,32,33,36,64,81–83^ and extensive experience with all experimental procedures.

### Surgical Preparation

Neurophysiology experiments were performed as described previously^25,26,33,84,85^. Rats were induced with 2.5-3.0% isoflurane in oxygen in a plexiglass chamber and transferred to a heated surgical table where anesthesia was continued (2.5-3.0% isoflurane; 60% oxygen, balance nitrogen). Body temperature was monitored via rectal thermometer (Fisher Scientific, Pittsburgh, PA) and maintained between 37.5 ± 1.0°C during experiments. A polyethylene catheter (i.d. 1.67 mm; PE 240; Intramedic, MD) was inserted into the trachea *via* mid-line neck incision, and rats were mechanically ventilated (0.07mL/10g bw/ breath; 72 breaths/min; VentElite small animal ventilator; Harvard Apparatus, Holliston, MA, USA). End-tidal PCO_2_ was monitored throughout the experiment with a breathe-through capnograph with sufficient response time to monitor true end-tidal PCO_2_ in rats (Capnogard, Novametrix, Wallingford CT). The inspired CO_2_ fraction was adjusted as needed to keep end-tidal PCO_2_ within pre-determined levels.

Anesthesia was slowly converted to urethane *via* slow infusion into a tail vein catheter (1.8 g/kg at 6mL/h; 24 gauge, Surflo, Elkton, MD), while progressively decreasing inspired isoflurane concentration 0.5% every 6 min. Anesthetic depth was verified by lack of physiological response to toe-pinch; supplemental anesthetic was infused if required. Rats were bilaterally vagotomized at the mid-cervical level to prevent phrenic nerve entrainment with the ventilator. The right femoral artery was isolated and cannulated with polyethylene tubing (i.d. 0.58 mm; PE 50; Intradermic, MD) to monitor blood pressure (TA-100 Transducer Amplifier, CWE, Inc.) and sample blood gases (PaO_2_, PaCO_2_, pH, base-excess and hemoglobin concentration) with a blood gas analyzer (ABL 90 Flex, Radiometer, Copenhagen, Denmark). Once the conversion to urethane anesthesia was complete, intravenous fluids were administered to maintain fluid and acid-base balance (1.5 mL/h; started ∼1 h after the beginning of surgery, or ∼20 min after conversion to urethane was complete^85^; 1:4 of 8.4% Na_2_CO_3_ in lactated Ringer’s solution). Rats were then given neuromuscular block with pancuronium bromide (2 mg/kg; Sigma-Aldrich, St. Louis, MO).

### Spinal Adenosine and Inosine Measurements

In separate rats (n = 5), a dorsal midline incision was made from the base of the skull to the fifth cervical segment. Muscle layers were retracted to expose C3–C5 vertebrae. A laminectomy was performed at the C4 vertebrae. Extracellular changes in adenosine concentration were measured *via* differential enzymatic detection using adenosine and inosine micro-biosensors (Zimmer-Peacock, UK) positioned ∼1 mm lateral to the spinal midline between C3 and C4 (∼1.5 mm depth from dorsal root entry zone) as previously described^33^. Coordinates were obtained from previous studies^86^. A longitudinal cut was made in the dura to enable micro-biosensor insertion ∼3–4 mm apart from each other on the same side of the spinal cord. Extracellular adenosine concentration was measured: 1) for 30 min following intrathecal Fkn protein (100 ng) delivery; or 2) during 5 min moderate (PaO_2_ = 40–55 mmHg) or severe (PaO_2_ = 25–30 mmHg) hypoxic episodes. Signals were acquired at 1 Hz and converted to concentrations *via* 3-point calibration. Measurements began ∼30–45 min after probe insertion. At the end of experiments, the adenosine biosensor was retracted from the spinal cord and immediately placed in a calibration chamber containing buffer solution. The signal obtained following exogenous adenosine delivery to the chamber (10 µM), measured in nanoamperes, was used to scale the adenosine signal (assuming linear response)^33,87^.

### Neurophysiological Experiments

#### Intrathecal drug delivery

A laminectomy was performed at the C_2_ vertebrae for intrathecal drug delivery. A small hole was cut in the dura near the junction of the C_2_ and C_3_ segments, and a flexible silicone catheter (O.D. 0.6 mm; Access Technologies) was advanced to the caudal end of C_3_. A 50 µL Hamilton syringe containing drug (*see* ‘Drugs and Vehicles’) was attached to the catheter for drug delivery. To prevent off target effects of pharmacological manipulation, we used intrathecal drug injections directly at C_4_ to limit unintended drug distribution. For example, in an anatomically separated motor pool (hypoglossal), mAIH-induced hypoglossal LTF is unaffected by C_4_ intrathecal drug delivery^26,27^, at least with injection volumes less than 20 µL.

#### Electrophysiological Recordings

Using a dorsal approach, the left phrenic nerve was isolated, cut distally and de-sheathed. Suction recording electrodes filled with 0.9% saline were placed in the saline-filled phrenic pocket and the nerve was suctioned with a syringe to record respiratory neural activity. Nerve activity was amplified (10K, A-M systems, Everett, WA), filtered (bandpass 300-5,000 Hz), integrated (time constant, 50 ms), digitized (CED 1401, Cambridge Electronic Design, UK), and analyzed using Spike2 software (CED, version 8.20). Inspiratory phrenic activity served as an index of respiratory motor output.

### Experimental Protocols

At least 45 min after full conversion to urethane anesthesia, the CO_2_ apneic and recruitment thresholds of phrenic nerve activity were determined by: 1) lowering inspired CO_2_ levels, or 2) increasing ventilation rate until rhythmic phrenic nerve activity ceased. After ∼1 min, inspired CO_2_ levels were slowly increased until rhythmic respiratory activity resumed. Baseline conditions were established by raising end-tidal PCO_2_ ∼2 mmHg above the recruitment threshold. After stable nerve activity was observed, blood samples were taken to document blood gas levels during baseline conditions. Arterial PCO_2_ was maintained isocapnic (± 1.5 mmHg) with respect to baseline blood gas values by actively manipulating inspired CO_2_ concentration and/or ventilation rate. Baseline oxygen levels (60% oxygen, balance nitrogen and carbon dioxide; PaO_2_ ≥ 150 mmHg) were maintained during experiments, except for hypoxic challenges (PaO_2_ = 40-55 mmHg). At the end of protocols, maximum chemoreflex activation was determined by delivering 7% CO_2_ and 10% O_2_ in N2 for ∼3 min to ensure preparation stability and to assure sufficient dynamic range in phrenic nerve burst amplitude (see *Supplement*). At the end of experiments, rats were euthanized by urethane overdose.

### Drugs and Vehicles

Drugs used include MSX-3 (A_2A_ receptor antagonist; #M3568; Millipore Sigma), Recombinant Rat CX3CL1 protein (#537-FT-025; FisherSci), AZD8797 (CX3CR1 antagonist; #2255; Axon Medchem), and ARL67156 (selective ecto-ATPase inhibitor; #128310; FisherSci). Upon arrival, all drugs were dissolved in 100% DMSO or 0.9% saline based on solubility and manufacturer recommendations. Aliquots of stock solutions were kept frozen at −20°C. On the day of experiments, drugs were diluted in sterile 0.9% saline to achieve the desired concentration. DMSO-saline ratios were determined by drug solubility; a final concentration of 10% DMSO was sufficient to dissolve all drugs in vehicle solution. Most drugs were dissolved to final effective concentrations previously determined in our laboratory *via* published dose-response studies^32^ or dose-response curves reported in the *Supplement*. Based on these (and other) reports, intrathecal drug doses were as follows: 10 µM MSX-3^33,77,88^, 100 ng Fractalkine protein, 10 µM AZD8797 and 1 mM ARL67156.

### Microglia Depletion

A group of rats was treated with Pexidartinib (PLX3397; MedKoo Biosciences, Morrisville, NC), a colony-stimulating factor 1 inhibitor. PLX3397 was formulated in DMSO, 1% PS80 and 0.5% hydroxycellulose. Rats were dosed at 80 mg/kg per day *via* syringe feeding (PLX3397 in DMSO formulation and liquefied Diet Gel; n=12) for 10 days. Rats were prepared for *in vivo* neurophysiology one day after the final dose.

### Phrenic Motor Neuron CtB and siRNA delivery

Intrapleural injections of the β-subunit of cholera toxin (CtB; Millipore Sigma, #C9903) were performed as described previously^46–48,73^. Fifteen μl of cholera toxin subunit B (30 μl total per animal, 2 μg μl^−1^ in sterile H_2_O) was injected intrapleurally on each side using 25 μl Hamilton syringes at a depth of 6 mm through the fifth intercostal space anterior axillary line 14 days prior to tissue harvest. After 11 days, a pool of small interfering RNAs (siRNA; Dharmacon, Inc.) targeting CX3CL1 (siFkn) mRNA (n=14) or a non-targeting sequence (siNTg; n=8) were delivered intrapleurally. Each siRNA consisted of 4 pooled 21-nucleotide duplexes with symmetrical 3’ overhangs (Accell SMARTpool). Target sequences of the 4 duplexes in the Fkn/CX3CL1 siRNA pool were:

1. UUAUCAACAUGAAUAGUACA
2. GGUUGGACUUUGUUGGUUC
3. CCAUUUUGUAUUUUACUAA
4. UUUUCAAGCAUCAUUACCA

Fkn and non-targeting siRNA (siNTg) were suspended in siRNA Universal Buffer (Dharmacon) to yield a concentration of 5 μM. The siRNA stocks were aliquoted and stored at −20 °C. Prior to injection, stock siRNA solutions (26.8 μl/injection; 100 pmol) were combined with Oligofectamine (3.2 μl/injection) and incubated at room temperature (22-24 °C) for 15 minutes.

Rats were lightly anesthetized with isoflurane in a closed chamber followed by inhalation of isoflurane *via* nose cone (2.5–3.5% in 50% O_2_, balance N_2_). Using aseptic technique and a sterile, RNase-free 25 μl Hamilton syringe, siRNAs were injected intrapleurally at the fifth intercostal space along the anterior axial line, as previously described^46–48^. Each rat received daily bilateral intrapleural injections of the appropriate siRNAs for 3 consecutive days. On the 4^th^ day, rats were prepared for *in vivo* neurophysiology experiments.

### In Situ Hybridization

Detection of Fkn/CX3CL1 mRNA using *in situ* hybridization was done to verify siRNA knockdown in phrenic motor neurons. On day 14, rats were anaesthetized under isoflurane anaesthesia (4% in O_2_) and transcardially perfused with ice-cold RNase-free 1X phosphate buffered solution (PBS) followed by 4% paraformaldehyde. Spinal segments C3-C5 were harvested and dehydrated in RNase-free 30% sucrose. Twenty micrometer sections were cut on a microtome and collected in 10-15 sets of serial C3-C5 sections in RNase-free PBS, in order to obtain representative staining across segments C3-C5. Sections were mounted on to Superfrost Plus Gold microscope slides (ThermoFisher Scientific, Inc., Waltham, MA). After drying at room temperature for ∼2 hours, slide-mounted tissue was stored at −80°C until use. All reagents used for *in situ* hybridization were purchased from Advanced Cell Diagnostics (Newark, CA). *In situ* hybridization was used for Fkn/CX3CL1 mRNA detection (reference no. 531141) using a fluorescence-based assay (reference no. 323110), as previously described^89^. Experimental protocols were performed according to the manufacturer instructions in *RNAscope Multiplex Fluorescent Detection Reagents V2* (Document No. UM 323100, Advanced Cell Diagnostics).

### Immunohistochemistry

After *in situ* hybridization experiments, immunohistochemistry for CtB and Iba-1 was done to detect labeling in phrenic motor neurons and activated microglia, respectively. Slides were rinsed 3 times in TBS, treated with blocking solution (10% donkey serum and 0.1% TritonX-100 in TBS) at room temperature for 1 hour, and then incubated in primary antibody, goat anti-CtB (1:1500; reference no. 227040, Millipore) and rabbit anti-Iba-1 (1:500; reference no. 019-19741, Wako) overnight at 4°C. The following day, slides were washed in TBS three times and incubated in donkey anti-goat secondary antibody (1:1500; AlexaFluor 488, Jackson ImmunoResearch, reference no. 705-545-147) and donkey anti-rabbit secondary antibody (1:1500; Cy5, Jackson ImmunoResearch, reference no. 711-175-152) for 2 hrs at room temperature. Slides were rinsed in TBS 3 times and coverslipped with ProLong Gold + DAPI antifade reagent (ThermoFisher Scientific). Immunohistochemistry for CtB and Iba-1 was similarly done following microglial depletion using PLX3397 to verify microglial knockdown in spinal C3-C6 ventral horn near the phrenic motor nucleus. Negative controls omitting either primary or secondary antibodies confirmed minimal nonspecific signal.

### Imaging and Acquisition

Slides were imaged using a fluorescence/brightfield microscope (BZ-X710, Keyence Co., Osaka, Japan) with a 20x or 40x lens. Fkn mRNA was detected using a TexasRed filter (model no: OP-87765); CtB immunolabelling was detected via a GFP filter (model no: OP-87763). Iba1 immunolabeling was detected via a Cy5 filter (model no. OP-87766). DAPI was detected via a DAPI filter (model no: OP-87762). The same light exposures were used for both groups (20X: 1/20 s, 1/25 s, 1/30 s and 1/200 s; 40X: 1/8.5 s, 1/3 s, 1/4 s and 1/12 s for Fkn mRNA, CtB, Iba1 and DAPI, respectively). Representative images were obtained with a confocal microscope (FV1000, Olympus Fluoview).

### Image Quantification

Fluorescence intensity was analyzed with QuPath Software (version 0.4.3) using images localized to the CtB-labeled area of the ventral horn. Background intensity level was determined using the median value of a control region (*i.e.,* near the central canal for each side). A threshold for each image was determined by constructing a pixel intensity histogram for each image and selecting a fixed percentile value across all images, as previously described^46,89^. The QuPath “wand” tool was used to annotate CtB-positive (phrenic motor neuron) and negative (non-phrenic) cells; cell nuclei were identified using DAPI. Fluorescence intensity of Fkn mRNA labeling was calculated as the average intensity of the CtB-positive cells in each section and normalized to control region values. Intensities of individual non-phrenic motor neurons were calculated as previously described^46,89^. Fkn labeling in non-phrenic cervical motor neurons was measured to assess potential non-specific effects.

Image analysis was performed using MATLAB R2021b (Mathworks) to count microglia labeled with the fluorescent marker CY5 in images of rat spinal cord slices. The code reads in overlay images showing CY5-labeled microglia, performs background subtraction and thresholding to identify microglia somata, and counts the number of microglia in each image. Key parameters used in the analysis include: 1) image resolution (0.75488 μm/pixel); 2) radius for selecting microglia somata (35 μm); 3) percentile threshold for CY5 intensity (99.0); and 4) expected soma size (100 pixels).

Data across all measured cells of all sections were averaged for each rat, representing the measured value for that animal. All imaging and image analysis was done by blinded assessors.

### Statistical Analyses

Measurements of peak integrated phrenic burst amplitude and burst frequency (bursts/min) were assessed in 1 min bins immediately before each arterial blood sample at: baseline, during the last minute of the first hypoxic episode, 30, 60 and 90 min post-AIH, and during the final minute of the maximum chemoreflex challenge (***Supplementary Table 3***). Measurements were made at equivalent time points in time-matched control experiments. Integrated nerve burst amplitude was normalized by subtracting the baseline value, dividing the difference by the baseline value, and reporting percentage changes from baseline. Burst frequencies were also normalized to baseline, expressed as an absolute difference in bursts per minute. All statistical comparisons between treatment groups for nerve burst amplitude (baseline and 90 min) were made using a 2- or 3-way ANOVAs with a repeated measures design. Individual comparisons were made using the Tukey *post-hoc* test.

Comparisons of mean arterial pressures and arterial PCO_2_ and PO_2_ (***Supplementary Tables 1*** and ***2***) were made at baseline, during hypoxia episode 1, 30, 60 and 90 min post-AIH using a 2-way mixed effects ANOVA to test if there was an effect of protocol with drug pretreatment. Unpaired *t*-tests were used to compare measurements of adenosine concentration during moderate (PaO_2_ = 40-50 mmHg) *vs* severe (PaO_2_ = 25-30 mmHg) hypoxic episodes.

Fkn/CX3CL1 mRNA *in situ* hybridization and microglia counts were compared using unpaired *t*-test. All statistics were analyzed in SigmaPlot v.12.0. Differences between groups were considered significant if p < 0.050. Data are presented as mean ± standard error mean (SEM).

## DATA AVAILABILITY

The datasets generated during and/or analyzed during the current study are available from the corresponding author on reasonable request.

## ACKNOWLEDGEMENTS

The authors would like to thank Dr. Jordan Follet and Dr. Matthew Farrer’s Laboratory of Neurogenetics and Neuroscience (LNN) at the University of Florida for use of and assistance with the confocal microscope. Funding was provided by the National Institutes of Health grants R01HL149800 (GSM), R01HL148030 (GSM), T32HL134621-5 (ABM) and the Francis Family Foundation (AT). The funders had no role in study design, data collection and analysis, decision to publish or preparation of the manuscript. Figures created with BioRender.com.

## AUTHOR INFORMATION

These authors contributed equally: A.B. Marciante and A. Tadjalli. A.B.M., A.T., J.J.W., T.L.B., and G.S.M. conceived and designed research. A.B.M., A.T., K.A.B., J.O. and Y.B.S. performed experiments. A.B.M., A.T., and G.S.M. analyzed data. E.K.L. wrote and executed code for microglia cell counts. A.B.M., A.T., and G.S.M. interpreted results of experiments. A.B.M. prepared figures. A.B.M. and G.S.M. drafted manuscript. A.B.M., A.T., Y.B.S., M.N., J.J.W., and G.S.M. edited and revised manuscript. All authors approved the final version of this manuscript.

## ETHICS DECLARATIONS

The authors declare no competing interests.

