## Supplementary Data for "Microglia regulate motor neuron plasticity via reciprocal fractalkine/adenosine signaling"

**This PDF file includes:**

**Supplemental Figure 1**
**Supplemental Figure 2**
**Supplemental Figure 3**
**Supplemental Figure 4**

**Supplemental Table 1**
**Supplemental Table 2**
**Supplemental Table 3**

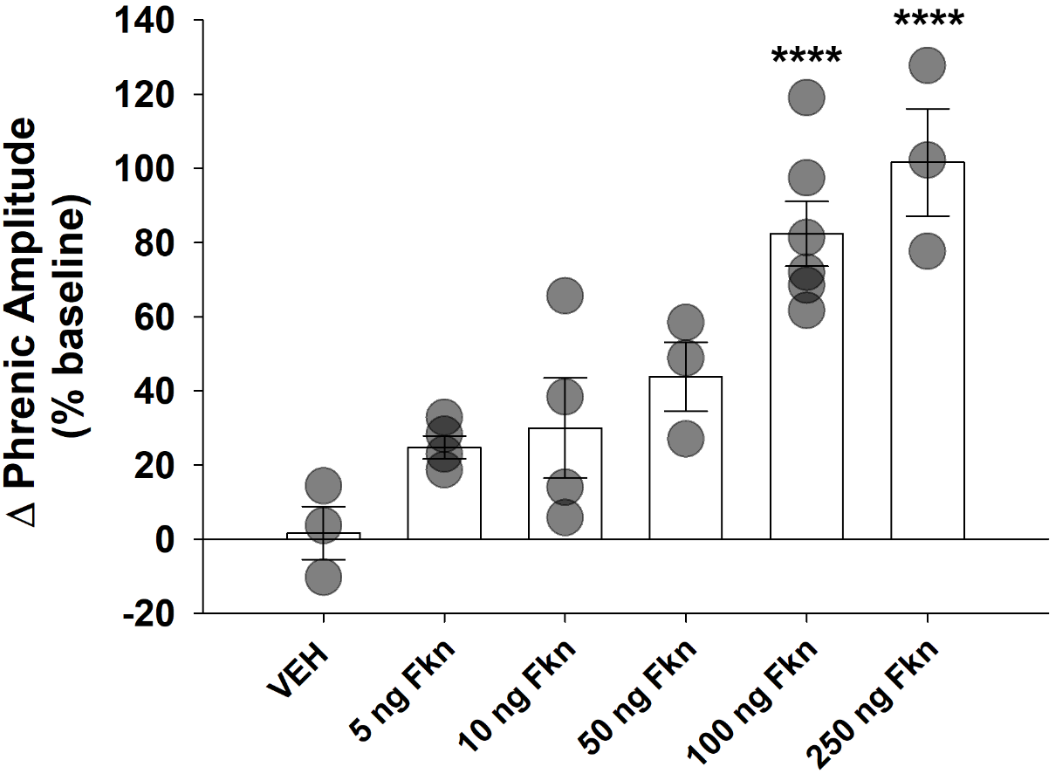


**Supplementary Figure 1.** ***Dose-response for intrathecal Fkn-induced phrenic motor facilitation.*** Intrathecal fractalkine (Fkn) or vehicle (VEH) was delivered at cervical spinal segment 4 at a concentration of 0, 5, 10, 50, 100 or 250 ng Fkn protein (CX3CL1). At 90 min post-administration, phrenic nerve amplitude was measured and normalized as a % change from baseline. Results are presented as mean ± SEM (n=3-6 independent experiments per group). Fkn concentration had a significant effect on the magnitude of phrenic motor facilitation (pMF; $F(5,17) = 13.041$, $p < 0.001$; One-Way ANOVA). pMF was significant *versus* VEH at 100 ng ($p < 0.001$; Tukey *post-hoc* Test) and 250 ng ($p < 0.001$; Tukey *post-hoc* Test) Fkn doses. ****, $p < 0.001$.

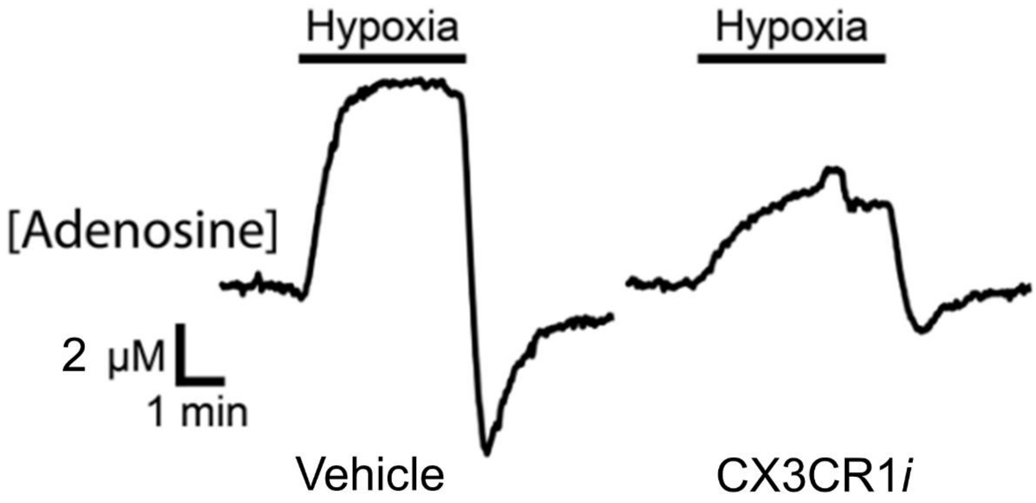


**Supplementary Figure 2.** ***Spinal fractalkine receptor inhibition suppresses spinal adenosine accumulation during hypoxic episodes.*** Intrathecal fractalkine receptor inhibitor (CX3CR*i*) or vehicle was delivered at cervical spinal segment 4. Average traces (n=3 recordings from 2 rats) for extracellular spinal adenosine concentration immediately prior to, during, and 5 min after severe hypoxia ($PaO_2$ = 28.8 ± 1.3 mmHg). CX3CR1 inhibition lowered peak and total adenosine concentration (area under curve) to ~ 1/3 of that observed in rats treated with vehicle.

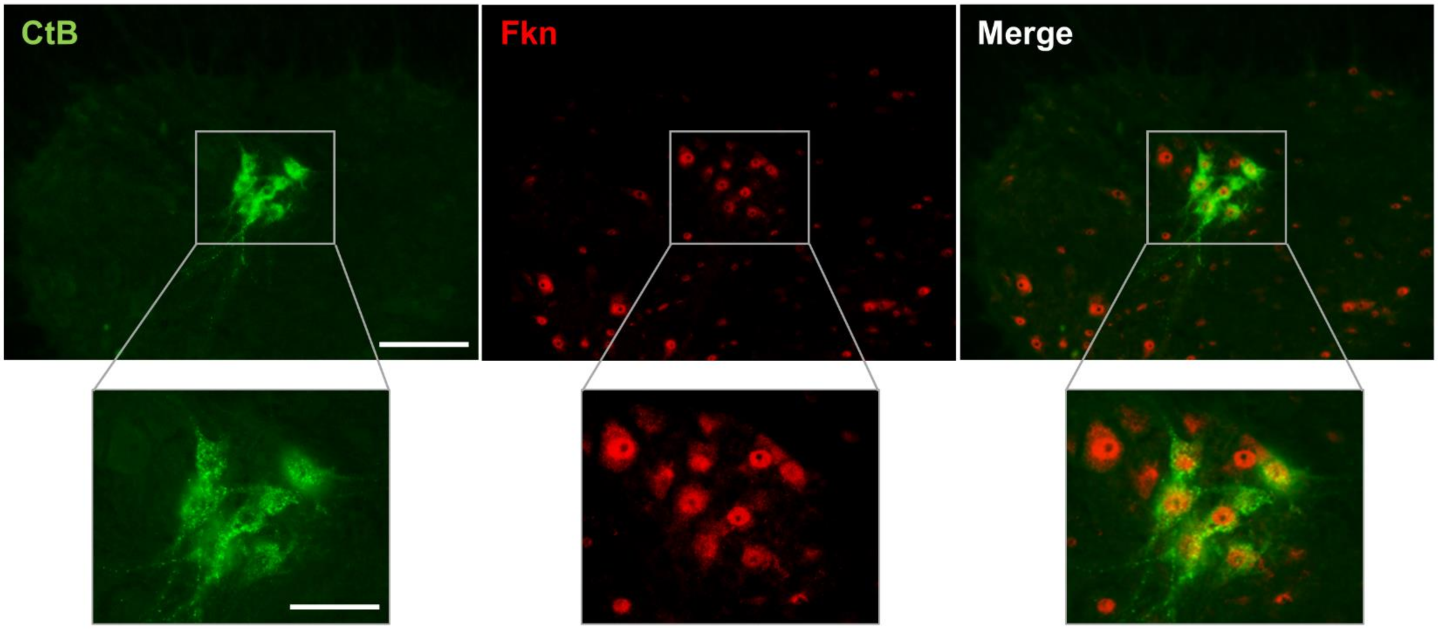


**Supplementary Figure 3.** ***Ventral horn phrenic and non-phrenic motor neurons express Fkn.*** Intrapleural Cholera toxin β (CtB) injections were administered to retrogradely-label phrenic motor neurons. Cervical spinal cord segments 3-6 were immunostained for CtB (green) and Fractalkine (Fkn; red). Representative images obtained from 6 individual rats. Scale bar (top row): 100 µm; Scale bar (bottom row): 50 µm.

| *Groups* | | *$Pa_{CO_2}$, mmHg* | | | | *$Pa_{O_2}$, mmHg* | | | | *MAP, mmHg* | | | |
|---|---|---|---|---|---|---|---|---|---|---|---|---|---|
| | n's | Baseline | 30 min | 60 min | 90 min | Baseline | 30 min | 60 min | 90 min | Baseline | 30 min | 60 min | 90 min |
| ***pMF Studies*** | | | | | | | | | | | | | |
| Vehicle Time Control | 7 | 42.3 ± 0.7 | 42.1 ± 0.8 | 42.7 ± 0.9 | 42.1 ± 0.9 | 293 ± 12 | 282 ± 6 | 282 ± 6 | 283 ± 9 | 113 ± 10 | 117 ± 5 | 119 ± 5 | 118 ± 4 |
| Vehicle + Fkn | 6 | 41.8 ± 1.6 | 41.6 ± 1.7 | 41.7 ± 1.8 | 42.8 ± 1.6 | 322 ± 7 | 304 ± 5 | 296 ± 7 | 290 ± 9[a] | 135 ± 6 | 124 ± 6 | 132 ± 5 | 131 ± 7 |
| AZD8797 + Fkn | 7 | 43.9 ± 1.8 | 43.6 ± 1.7 | 45.0 ± 2.2 | 43.9 ± 1.4 | 314 ± 17 | 297 ± 18 | 286 ± 18[a] | 274 ± 18[a] | 130 ± 4 | 124 ± 3 | 127 ± 3 | 130 ± 4 |
| ARL67156 + Fkn | 7 | 43.0 ± 2.2 | 43.1 ± 2.2 | 43.5 ± 2.1 | 43.1 ± 2.5 | 291 ± 11 | 280 ± 15 | 256 ± 17[a] | 252 ± 22[a] | 133 ± 7 | 130 ± 8 | 135 ± 9 | 137 ± 8 |
| MSX-3 + Fkn | 7 | 42.5 ± 1.0 | 41.8 ± 1.6 | 41.2 ± 1.2 | 43.0 ± 1.2 | 315 ± 11 | 298 ± 9 | 292 ± 9 | 294 ± 19 | 125 ± 8 | 117 ± 6 | 119 ± 6 | 114 ± 4 |
| ***Drug + pLTF Studies*** | | | | | | | | | | | | | |
| Vehicle + mAIH | 7 | 42.9 ± 1.3 | 43.1 ± 1.6 | 43.3 ± 1.3 | 43.3 ± 1.3 | 318 ± 10 | 251 ± 11[a] | 276 ± 6[a] | 279 ± 8 | 135 ± 7 | 115 ± 10[a] | 126 ± 14 | 125 ± 15 |
| Vehicle + sAIH | 7 | 43.7 ± 1.7 | 44.4 ± 1.5 | 43.5 ± 1.6 | 43.8 ± 1.6 | 312 ± 6 | 247 ± 12[a] | 266 ± 8[a] | 260 ± 10[a] | 137 ± 10 | 133 ± 13 | 129 ± 14 | 129 ± 16 |
| AZD8797 + mAIH | 7 | 44.1 ± 0.8 | 44.1 ± 1.0 | 43.8 ± 1.1 | 44.3 ± 1.2 | 312 ± 6 | 247 ± 12[a] | 266 ± 8[a] | 260 ± 10 | 133 ± 5 | 129 ± 6 | 116 ± 8[a] | 120 ± 5 |
| AZD8797 + sAIH | 7 | 44.9 ± 1.0 | 45.2 ± 1.0 | 45.4 ± 1.0 | 45.1 ± 1.1 | 328 ± 5 | 258 ± 15[a] | 277 ± 8[a] | 279 ± 9[a] | 114 ± 2 | 113 ± 5 | 115 ± 6 | 115 ± 7 |
| PLX3397 + mAIH | 4 | 41.4 ± 0.7 | 42.9 ± 1.2 | 42.5 ± 1.3 | 42.1 ± 0.6 | 334 ± 6 | 213 ± 18[a,c] | 238 ± 8[a] | 267 ± 6[a] | 136 ± 4 | 132 ± 1[b] | 135 ± 1 | 136 ± 2 |
| PLX3397 + sAIH | 6 | 41.2 ± 0.5 | 42.0 ± 0.6 | 41.4 ± 0.6 | 41.6 ± 0.8 | 328 ± 5 | 258 ± 15[a] | 277 ± 8[a] | 279 ± 9[a] | 138 ± 7 | 134 ± 5 | 133 ± 6 | 133 ± 4 |
| Fkn + mAIH | 8 | 42.7 ± 0.8 | 42.8 ± 1.6 | 42.2 ± 1.2 | 42.9 ± 1.0 | 313 ± 10 | 203 ± 15[a] | 228 ± 15[a] | 240 ± 13 | 127 ± 6 | 123 ± 8 | 138 ± 4 | 137 ± 5 |
| Fkn + sAIH | 5 | 41.5 ± 1.5 | 42.6 ± 1.9 | 40.6 ± 1.4 | 41.1 ± 1.5 | 304 ± 18 | 203 ± 38[a] | 273 ± 16 | 268 ± 10 | 129 ± 9 | 123 ± 12 | 124 ± 13 | 124 ± 15[a] |
| ***RNAi + pLTF Studies*** | | | | | | | | | | | | | |
| siNontargeting + mAIH | 4 | 41.7 ± 2.7 | 41.5 ± 2.7 | 42.2 ± 2.3 | 42.5 ± 2.3 | 308 ± 5 | 219 ± 34[a] | 253 ± 40 | 270 ± 44 | 133 ± 2 | 129 ± 8 | 129 ± 9 | 129 ± 8 |
| siNontargeting + sAIH | 4 | 43.0 ± 1.6 | 43.2 ± 1.8 | 43.6 ± 1.5 | 43.8 ± 2.1 | 348± 21 | 206 ± 37[a,c] | 293 ± 23 | 314 ± 14 | 139 ± 11 | 134 ± 6 | 133 ± 8 | 135 ± 11 |
| siFkn + mAIH | 7 | 40.7 ± 0.5 | 42.1 ± 0.4 | 41.9 ± 0.6 | 41.2 ± 0.3 | 348 ± 8 | 208 ± 25[a,c] | 319 ± 20 | 305 ± 21 | 139 ± 5 | 135 ± 6 | 139 ± 6 | 141 ± 6 |
| siFkn + sAIH | 7 | 43.3 ± 0.5 | 43.6 ± 0.6 | 43.8 ± 0.6 | 44.0 ± 0.6 | 317 ± 15 | 243 ± 38[a] | 280 ± 12 | 276 ± 27 | 129 ± 9 | 133 ± 3 | 132 ± 4 | 134 ± 5 |

$p < 0.050$ indicates a significant difference for:

[a], different from baseline
[b], different from appropriate time control
[c], different from 90 min measurement

**Supplementary Table 1.** *Arterial $P_{CO_2}$, $P_{O_2}$, and MAP during baseline and 30, 60 and 90 min post-AIH.*

| *Groups* | *Hypoxia* | | |
|---|---|---|---|
| | n's | *$PaCO_2$, mmHg* | *$PaO_2$, mmHg* |
| ***Drug + pLTF Studies*** | | | |
| Vehicle + mAIH | 7 | 41.5 ± 1.4 | 40.7 ± 4.5[a] |
| Vehicle + sAIH | 7 | 43.1 ± 1.8 | 25.6 ± 0.6[a,b] |
| AZD8797 + mAIH | 7 | 42.9 ± 1.3 | 43.3 ± 1.9[a] |
| AZD8797 + sAIH | 7 | 44.2 ± 1.7 | 26.8 ± 0.8[a, b] |
| PLX3397 + mAIH | 4 | 42.2 ± 0.5 | 45.3 ± 2.4[a] |
| PLX3397 + sAIH | 6 | 41.3 ± 0.6 | 27.3 ± 0.3[a, b] |
| Fkn + mAIH | 7 | 41.6 ± 1.1 | 44.4 ± 1.0[a] |
| Fkn + sAIH | 4 | 41.2 ± 1.8 | 28.4 ± 1.3[a, b] |
| ***RNAi + pLTF Studies*** | | | |
| siNontargeting + mAIH | 4 | 42.0 ± 3.1 | 41.2 ± 1.4[a] |
| siNontargeting + sAIH | 4 | 42.3 ± 2.1 | 28.4 ± 1.2[a, b] |
| siFkn + mAIH | 7 | 40.5 ± 0.9 | 46.5 ± 1.4[a] |
| siFkn + sAIH | 7 | 41.8 ± 0.5 | 26.3 ± 1.5[a, b] |

$p < 0.050$ indicates a significant difference for:

[a], different from baseline

[b], different from mAIH protocol with the same drug

**Supplementary Table 2.** *Arterial $P_{CO_2}$, $P_{O_2}$, and MAP during hypoxic episodes*

| *Groups* | *Baseline Nerve Amplitude (V)* | *Maximum Nerve Amplitude (V)* |
|---|---|---|
| ***pMF Studies*** | | |
| Vehicle Time Control | 0.120 ± 0.021 | 0.244 ± 0.040[a] |
| Vehicle + Fkn | 0.101 ± 0.009 | 0.234 ± 0.023[a] |
| AZD8797 + Fkn | 0.109 ± 0.010 | 0.265 ± 0.021[a] |
| MSX-3 + Fkn | 0.087 ± 0.009 | 0.220 ± 0.020[a] |
| ARL67156 + Fkn | 0.097 ± 0.021 | 0.241 ± 0.049[a] |
| ***Drug + pLTF Studies*** | | |
| Vehicle + mAIH | 0.074 ± 0.021 | 0.253 ± 0.024[a] |
| Vehicle + sAIH | 0.075 ± 0.014 | 0.234 ± 0.032[a] |
| AZD8797 + mAIH | 0.074 ± 0.007 | 0.254 ± 0.023[a] |
| AZD8797 + sAIH | 0.072 ± 0.012 | 0.267 ± 0.030[a] |
| PLX3397 + mAIH | 0.062 ± 0.015 | 0.196 ± 0.031[a] |
| PLX3397 + sAIH | 0.081 ± 0.023 | 0.238 ± 0.048[a] |
| Fkn + mAIH | 0.073 ± 0.011 | 0.159 ± 0.023[a] |
| Fkn + sAIH | 0.078 ± 0.015 | 0.205 ± 0.035[a] |
| ***RNAi + pLTF Studies*** | | |
| siNontargeting + mAIH | 0.058 ± 0.023 | 0.175 ± 0.058[a] |
| siNontargeting + sAIH | 0.048 ± 0.015 | 0.195 ± 0.047[a] |
| siFkn + mAIH | 0.040 ± 0.006 | 0.171 ± 0.018[a] |
| siFkn + sAIH | 0.056 ± 0.013 | 0.143 ± 0.035[a] |

$p < 0.050$ indicates a significant difference for:

[a], different from baseline

**Supplementary Table 3.** *Phrenic nerve amplitude, in volts (V), at baseline and during maximum chemoreceptor stimulation*
